# Comparative performance of mutual information and transfer entropy for analyzing the balance of information flow and energy consumption at synapses

**DOI:** 10.1101/2020.06.01.127399

**Authors:** Mireille Conrad, Renaud B Jolivet

**Affiliations:** Département de Physique Nucléaire et Corpusculaire, University of Geneva, Geneva, Switzerland

**Keywords:** Information theory, Mutual information, Transfer entropy, Synapse, Energetic optimality

## Abstract

Information theory has become an essential tool of modern neuroscience. It can however be difficult to apply in experimental contexts when acquisition of very large datasets is prohibitive. Here, we compare the relative performance of two information theoretic measures, mutual information and transfer entropy, for the analysis of information flow and energetic consumption at synapses. We show that transfer entropy outperforms mutual information in terms of reliability of estimates for small datasets. However, we also show that a detailed understanding of the underlying neuronal biophysics is essential for properly interpreting the results obtained with transfer entropy. We conclude that when time and experimental conditions permit, mutual information might provide an easier to interpret alternative. Finally, we apply both measures to the study of energetic optimality of information flow at thalamic relay synapses in the visual pathway. We show that both measures recapitulate the experimental finding that these synapses are tuned to optimally balance information flowing through them with the energetic consumption associated with that synaptic and neuronal activity. Our results highlight the importance of conducting systematic computational studies prior to applying information theoretic tools to experimental data.

**Author summary:** Information theory has become an essential tool of modern neuroscience. It is being routinely used to evaluate how much information flows from external stimuli to various brain regions or individual neurons. It is also used to evaluate how information flows between brain regions, between neurons, across synapses, or in neural networks. Information theory offers multiple measures to do that. Two of the most popular are mutual information and transfer entropy. While these measures are related to each other, they differ in one important aspect: transfer entropy reports a directional flow of information, as mutual information does not. Here, we proceed to a systematic evaluation of their respective performances and trade-offs from the perspective of an experimentalist looking to apply these measures to binarized spike trains. We show that transfer entropy might be a better choice than mutual information when time for experimental data collection is limited, as it appears less affected by systematic biases induced by a relative lack of data. Transmission delays and integration properties of the output neuron can however complicate this picture, and we provide an example of the effect this has on both measures. We conclude that when time and experimental conditions permit, mutual information – especially when estimated using a method referred to as the ‘direct’ method – might provide an easier to interpret alternative. Finally, we apply both measures in the biophysical context of evaluating the energetic optimality of information flow at thalamic relay synapses in the visual pathway. We show that both measures capture the original experimental finding that those synapses are tuned to optimally balance information flowing through them with the concomitant energetic consumption associated with that synaptic and neuronal activity.

## Introduction

The brain is commonly thought of as an information transmission, processing and storage biological machine, calling for comparisons to man-made devices with similar functions such as computers (see [1] for a discussion of these issues). As a consequence, tools and methods pertaining to such devices have been ported to the neurosciences for the study of neural networks. One such tool is information theory, designed in the late 1940s by Claude Shannon to formalize and find limits on signal processing and communication operations in machines [2].

Information theory has a long and successful history in neuroscience, where it has been applied to a variety of experimental data and theoretical contexts, and to address a variety of questions. One area of particular interest is the application of information theoretic concepts to spike trains, as they easily lend themselves to a reduction to binary sequences, whereupon action potentials are converted to 1s and the rest of electrophysiological traces to 0s. This reduction of spike trains to binary sequences has been used to measure information flow through synapses and neural networks, and propagation of information from the environment to the cortex through sensory pathways for instance [3-10]. The information theoretic quantity most often encountered in such contexts is the mutual information (*I*) [4]. The mutual information of two random variables – for instance two spike trains, or a sensory signal and a response spike train – is a measure of the mutual dependence between those two variables. By construction, *I* is symmetrical and quantifies how much information can be obtained about one random variable by observing the other one.

In 1998, Strong and colleagues published a procedure to measure *I* between a sensory input signal and the corresponding response of a neuron [11]. Through clever design, their so-called ‘direct method’, allows calculating *I* by measuring only the activity of the neuron of interest in response to a specifically designed input sequence containing repeating portions. This method has been successfully applied in a number of contexts, for instance to quantify information flow from the retina to the primary visual cortex in primates [3], or to quantify the relation between information flow and energy consumption at synapses in the visual pathway [8]. One limitation of using *I* is the need to collect relatively large amounts of data to avoid systematic evaluation biases, and a number of methods have been devised to compensate those biases when data is limited or difficult to acquire [6]. This, however, can be tricky in an experimental context, as the time available for collecting data can be limited for a large number of various reasons.

Mutual information does not however strictly quantify directional information flow as it is symmetrical by design. To address this limitation, Schreiber has proposed a modified version of mutual information called transfer entropy (*TE*), which is explicitly built to measure how much information flows from one random variable to another one, and which is thus not symmetrical [12]. While *TE* is used widely outside of neuroscience and in systems neuroscience, it is not used very often as a replacement of *I* for analyzing spike trains specifically.

We have recently published a series of experimental and computational works on the trade-off between information flow and concomitant energy consumption in neurons and neural networks, in which we used either *I* [7-9] or *TE* [10]. This work is part of an emergent interest for energetic questions in neural information processing [13-16]. Here, we proceed to a systematic comparison of both those quantities in different biologically-relevant scenarios when comparing inputs to a synapse and the output spike train generated by the postsynaptic neuron in response, using spike train data generated from Poisson processes, or an experimentally-calibrated biophysical model of thalamic relay cells of the Hodgkin-Huxley-type [8]. We decided to focus on those two measures, and specifically on the so-called direct method to compute *I*, because of their relative user-friendliness, which should make them popular methods among experimentalists. We report that while *TE* and *I* both allow accurate and consistent predictions of theoretical expectations in simple scenarios, *TE* is far less biased than *I* when little data is available and might thus offer more accurate measurements in experimental conditions where acquisition of large datasets is impossible or prohibitively costly. When used to analyze more realistic synthetic data (generated by a biophysical model of the Hodgkin-Huxley-type) however, the measure of *TE* can be strongly affected by systematic time frame shits between inputs and outputs, a problem *I* is immune to by construction when using the direct method. Finally, we show how both measures perform when applied to assessing the energetic optimality of information transfer at biophysically-realistic synapses, and compare those results to results from the literature.

Our results illustrate the importance of systematically testing information theoretic measures on synthetic test data prior to designing experiments in order to fully understand how much data needs to be collected, and understand the trade-offs involved in using different measures. Our results also provide some guidance on what measure (*I* or *TE*) will perform best under different circumstances.

## Results

In order to compare the relative performances of mutual information (*I*) and transfer entropy (*TE*) on measuring information flow at synapses, i.e. when comparing the input to a synapse to the output generated by the postsynaptic neuron, we started by generating synthetic binary input and output spike trains. To mimic the transmission of action potentials at thalamic relay cells, which we have studied and modelled before [8], we generated as input random binary Poisson spike trains at a fixed frequency matching what we had observed experimentally. That input sequence was then copied to form the output sequence with deletions occurring with a certain probability (non-transmitted spikes), and with additions (to mimic spontaneous firing of the output neuron). Numerical values derived from previous experiments for these simulations are given in Table 1 below. We have also previously studied transmission of information at the cortical synapse between thalamic relay cells and layer 4 spiny stellate cells [10]. We thus generated a second additional set of simulations using parameters matching the experimental observations for that second scenario. Numerical values for those simulations are also given in Table 1. Note that in both of these scenarios, information flows unidirectionally in a feed-forward manner (see [8] and [10] for further details).

**Table 1.** Parameters for generation of inputs and outputs of random spike trains.

| Scenario | Input frequency (Hz) | Transmission failure probability (per input spike) | Spontaneous firing probability (per output time bin) |
| --- | --- | --- | --- |
| Thalamic relay synapse | 20 ¶ | 0.8 ¶ | 0.00084 ¶ |
| Cortical layer 4 spiny stellate cell synapse | 4 § | 0.9 § | 0.0024 § |

¶ From ref. [8].
§ From ref. [10].

To compute the mutual information *I* between those input and output sequences, we used the so-called direct method. The direct method requires repeating sequences (see Methods), while this isn’t necessary for computing the transfer entropy (*TE*). We thus generated in each case two datasets, one with repetitions to use with the direct method to compute *I* (Figure 1A), and one without repetitions to compute *TE* (Figure 1B). In each case, the two datasets had the same duration, with the duration of the repeating sequence multiplied by the number of repetitions used to compute *I* equating the total duration of the dataset generated to compute *TE*. Unless mentioned otherwise, the time bin is always 3 ms in duration, approximately the temporal extension of an individual action potential.

**Figure 1.**
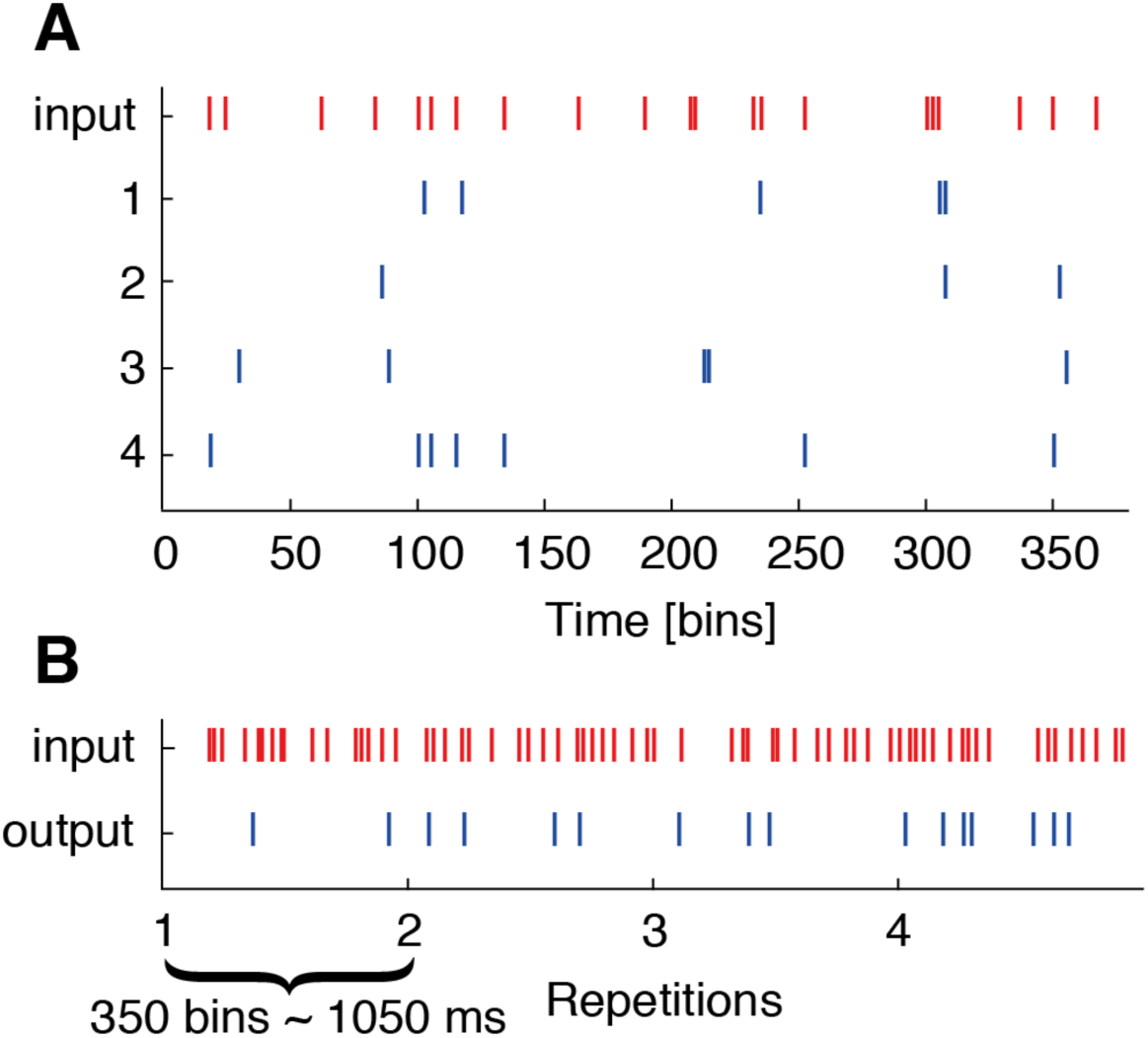
Sample input and output sequences for computing *I* and *TE*. Examples of input and output data generated, with repetitions for mutual information calculations (*I*; to use with the direct method) (**A**), and without repetitions for transfer entropy calculations (*TE*) (**B**). Here, outputs are generated using transmission statistics derived from experiments in the thalamic relay cell scenario (see Methods and Table 1). In each case, two length-matched datasets are generated to compare the relative performances of *I* and *TE* for recordings of a certain duration. 1 time bin = 3 ms.

The mutual information (*I*) can be defined as the difference between the two entropies *H*_*total*_ and *H*_*noise*_ (*I = H*_*total*_ *-H*_*noise*_; see Methods), and these entropies are typically calculated for ‘words’ of a certain length. For instance, the binary sequence 000110100100100101 can be segmented in words of length 1 yielding the words 0|0|0|1|1|…, or segmented in words of any other length. For words of length 3 for instance, it would yield 000|110|100|100|100|… In neuroscience, using long words is important to accurately capture and account for the information carried by stereotypical temporal patterns of spikes, if any [3, 11]. However, using longer and longer words can lead to significant biases in estimating *H*_*total*_ and *H*_*noise*_, and eventually *I*, when using a finite dataset to build estimates, which they are always. This can be a serious limitation when trying to use *I* in an experimental context where only limited data is available. To compensate for this, the so-called direct method includes two corrections when calculating *I*. The first correction factor is meant to extrapolate the true value of each individually computed entropies *H*_*total*_ and *H*_*noise*_ entering in the calculation of *I* to a dataset of infinite size, while the second correction is meant to extrapolate entropy estimates for both *H*_*total*_ and *H*_*noise*_ to infinitely long words. We have previously reported that these corrections did not lead to large changes in evaluating *I* when using limited experimental datasets with statistics similar to the datasets we generated here [8]. Additionally, we have had difficulties in reliably using the first above-mentioned correction (in [8] and here). As a consequence, here, we only implemented the second above-mentioned correction when using the direct method to compute *I*. Figure 2 shows how this correction was implemented. For *I, H*_*total*_ and *H*_*noise*_ were plotted as a function of the inverse of the word length and fitted with a linear function. We then extrapolated the fits to 0 (i.e. to words of infinite length) and took the difference of those values to calculate the mutual information (*I*). For transfer entropy, similar curves were computed, one for ‘raw transfer entropy’ and one for ‘noise transfer entropy’ (see Materials & Methods) [10, 17]. *TE* was then computed as the difference between those two curves, like for *I. TE* was then fitted with a linear relationship and extrapolated to infinite word lengths.

**Figure 2.**
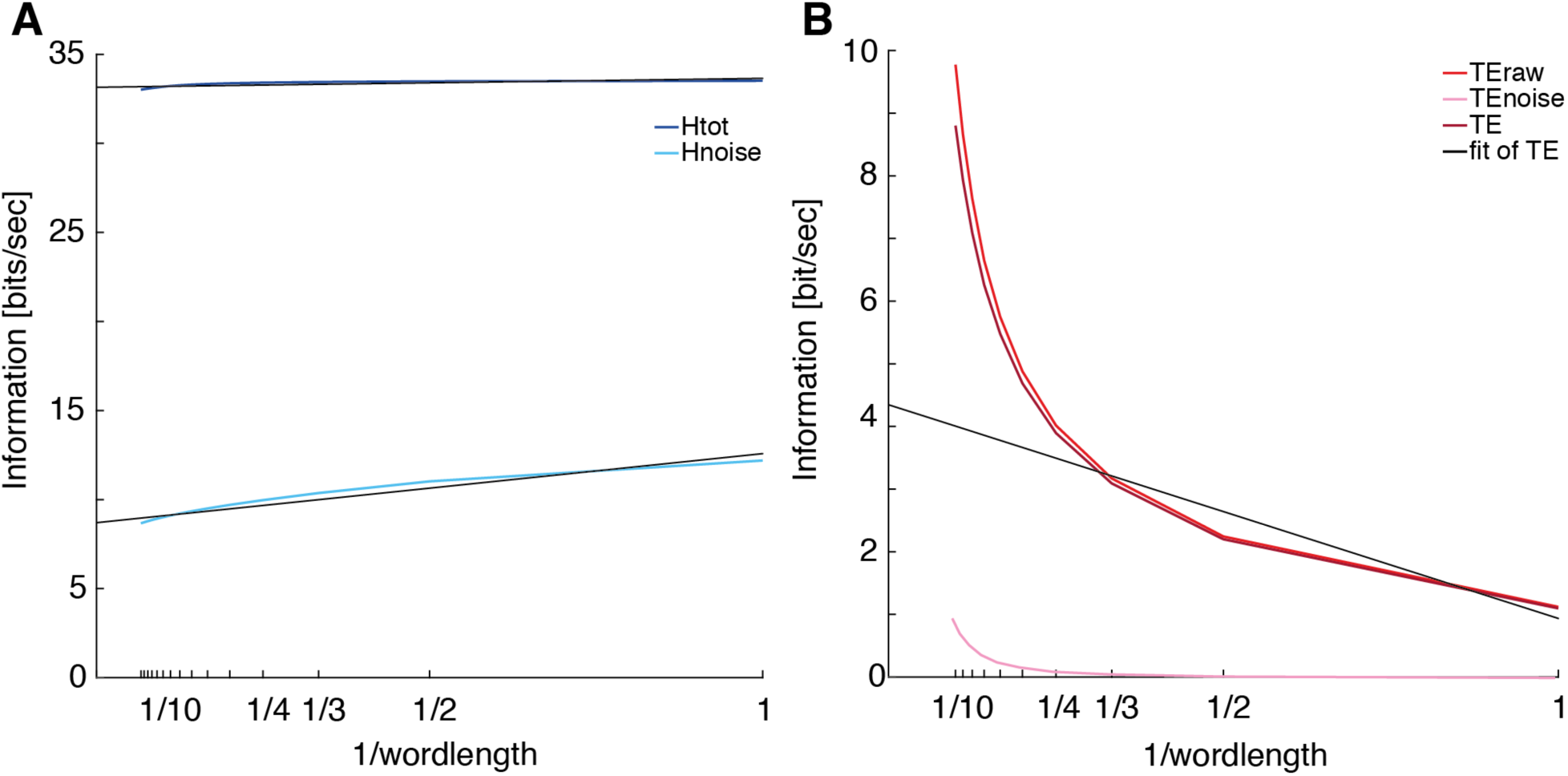
Extrapolation of mutual information and transfer entropy values to infinite word lengths. Example of extrapolation to infinite word lengths for mutual information (**A**) and transfer entropy calculations (**B**). Mutual information is calculated using the ‘direct’ method and the second correction of that method is applied to extrapolate both *H*_*tot*_ and *H*_*noise*_ to words of infinite lengths. The mutual information *I = H*_*tot*_ -*H*_*noise*_ at the intercept for 1/wordlength = 0. A similar method was used to evaluate the transfer entropy. In both cases, curves were generated using spike trains simulated using the Hodgkin-Huxley-type model.

In that relatively simple scenario of action potentials being transmitted at a single relay synapse with set probabilities, *TE = I*, and it is possible to calculate that value exactly (see Methods). Figure 3 shows a comparison between that theoretical value, and *I* and *TE* calculated as described above for parameters corresponding to action potential transmission at thalamic relay cells (A) [8] or layer 4 spiny stellate cells (B) [10] (see Table 1 for details) for datasets of increasing sizes, plotted as the equivalent number of repetitions as used to calculate *I* using the direct method. The calculated theoretical value for the thalamic relay cell scenario (Figure 3A, first line in Table 1 [8]) was 15.47 bits/sec. The calculated theoretical value for the layer 4 spiny stellate cell scenario (Figure 3B, second line in Table 1 [10]) was 1.54 bits/sec. Both these values closely match the experimental and theoretical results we had previously published. While it is apparent that both *TE* and *I* eventually converge to the expected theoretical value given a dataset of sufficient size, *TE* appears to perform vastly better for small datasets, converging faster (i.e. for smaller datasets) than *I* to the correct theoretical value. Furthermore, like *I, TE* overestimates the expected value of transmitted information, but as illustrated in the insets in Figure 3A and B, does so to a much lesser extent than *I*, even for very short datasets.

**Figure 3.**
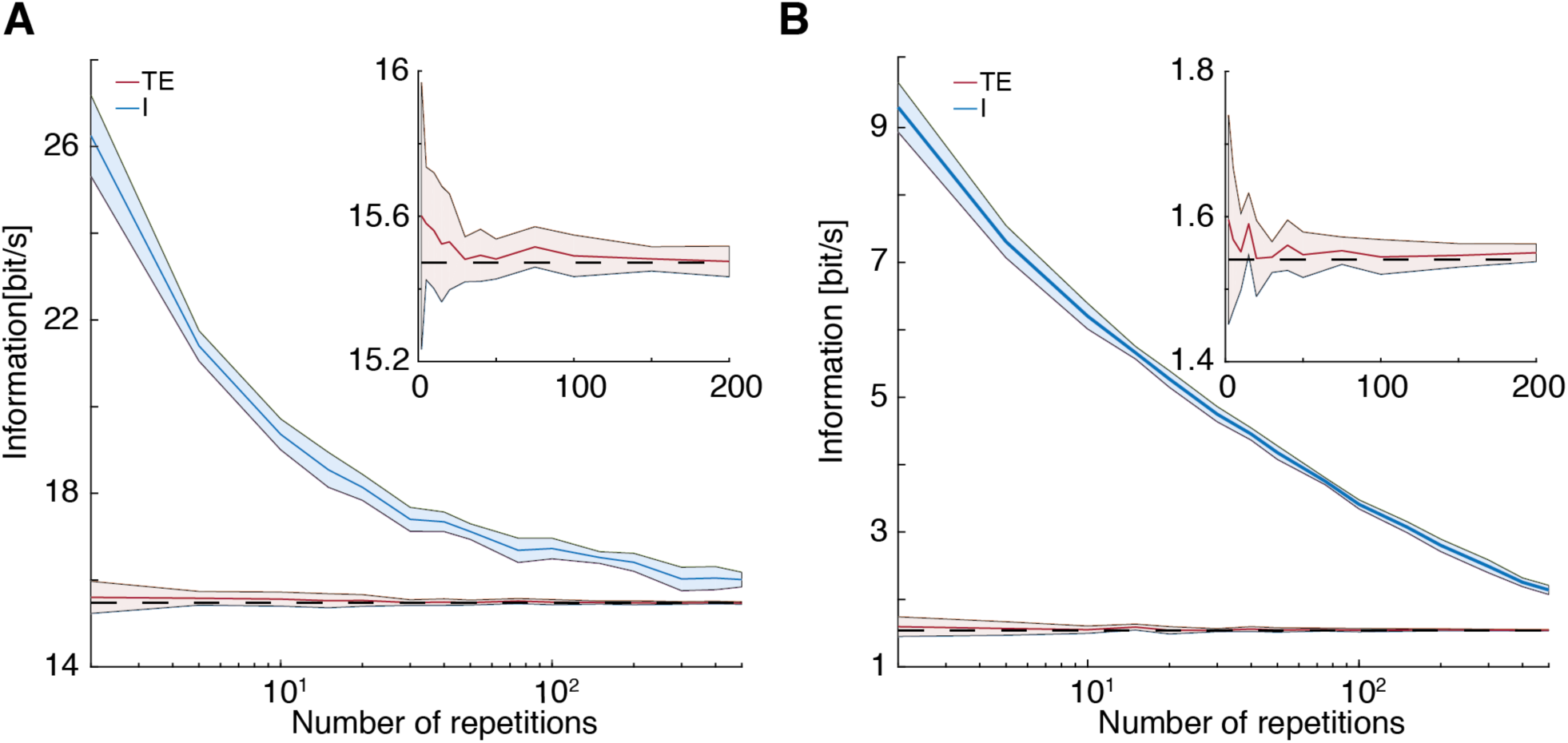
Comparative performance of *I* and *TE* in evaluating information flow at synapses driven by Poisson synthetic spike trains. Comparison between mutual information (*I*; blue; calculated following [8, 11]) and transfer entropy (*TE*; red; calculated following [10, 17]) as a function of the size of the dataset for randomly generated spike trains based on thalamic relay cells characteristics (**A**) [8] or layer 4 spiny stellate cells characteristics (**B**) [10] (see also Table 1). In each case, the black line indicates the theoretical value (see Methods). In both A and B, the inset zooms on *TE* for a low number of repetitions. In each case, shaded areas indicate the standard error of the mean.

Spike trains *in situ*, however, rarely display Poisson statistics. In the visual pathway for instance, spike trains impinging from the retina onto thalamic relay cells are characterized by non-Poissonian statistics with a high probability of spike doublets with short (∼10 ms) interspike intervals. In order to test if the results of Figure 3 depend on the temporal structure of the input spike train, we generated a second set of simulations as above, but replacing the Poisson input spike trains with spike trains generated using statistics matching *in situ* recordings. To do so, we computed the interspike interval distribution of input spike trains from experimental data collected in [8] and [10] (Figure 4A and B insets). We then used these to calculate the cumulative distribution function (CDF) of interspike intervals (Figure 4 insets). Finally, we used the cumulative distribution function to generate spike trains as a series of interspike intervals with a temporal structure matching what had been observed in experiments. Under these new circumstances, there is *a priori* no expectation that *TE* should be equal to *I*, and it is not possible to calculate the expected theoretical values simply. Figure 4 shows, however, that this does not significantly change the behavior of *TE* and *I* with respect to the size of the dataset used to compute them, with both *TE* and *I* converging to values very similar to the values in Figure 3 given a large enough dataset. Like in Figure 3, *TE* appears to significantly outperform *I* for short datasets. This prompts the question of why *TE* appears to systematically outperform *I* in the two relatively simple scenarios we tested here (Figures 3 and 4). In the Discussion below, we attempt to provide an answer to this question based on the structure of both measures.

**Figure 4.**
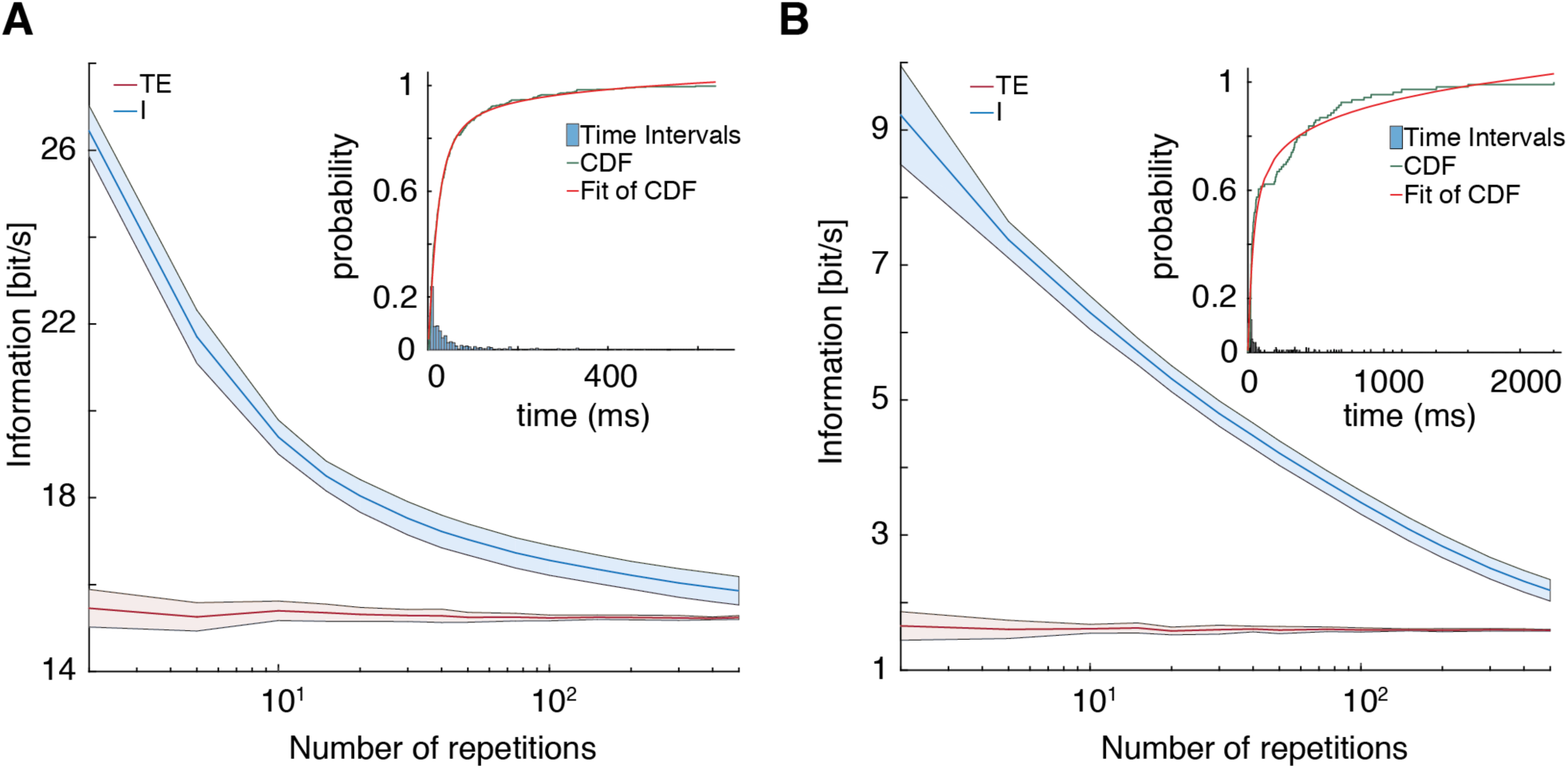
Comparative performance of *I* and *TE* in evaluating information flow at synapses driven by synthetic spike trains with realistic biological temporal structures. Comparison between mutual information (*I*; blue; calculated following [8, 11]) and transfer entropy (*TE*; red; calculated following [10, 17]) as a function of the size of the dataset for randomly generated using the cumulative distribution function (CDF) of the interspike intervals (Insets) based on experimental data recorded impinging onto thalamic relay cells (**A**) [8] or layer 4 spiny stellate cells (**B**) [10]. In each case, the inset shows the CDF of the biological interspike interval used to generate input sequences. In each case, shaded areas indicate the standard error of the mean.

We then wanted to test these comparisons using a more realistic model to generate output spike trains, focusing on the first scenario tested above: information transmission at thalamic relay synapses. To do so, we used an experimentally-calibrated biophysical single-compartment Hodgkin-Huxley-type model of thalamic relay cells (see [8] for details about the calibration of that model and Methods here for a detailed description of the model). To generate input for that model, we used the same procedure as described above for Figure 4, generating input spike trains with a temporal structure matching what is observed in *in vivo* experiments. We then convolved these spikes trains with unitary synaptic conductance extracted from experimental data in [8] (see Methods). Figure 5A shows a sample input spike train generated in this way, the corresponding synaptic conductance injected into the Hodgkin-Huxley-type model, the output spike train generated by that model in response to that synaptic input, and finally, the binarized output corresponding to that output spike train. We then applied *TE* and *I* on these binarized input and output spike trains in the same way as above. Figure 5C shows a plot comparing *TE* and *I* for datasets of increasing sizes (similar to what we plotted above in Figures 3A and 4A). It is immediately apparent that while the qualitative behavior of *I* and *TE* is not strikingly different than what we observed for spike trains generated using simple spiking and transmission probabilities as above, they do not appear to converge to similar values like they did previously (Figures 3A and 4A). *TE*, in particular, appears to converge to a value far lower than what we had previously observed (compare for instance Figure 5A to Figure 4A; the values reached by *I* in these two panels are more comparable).

**Figure 5.**
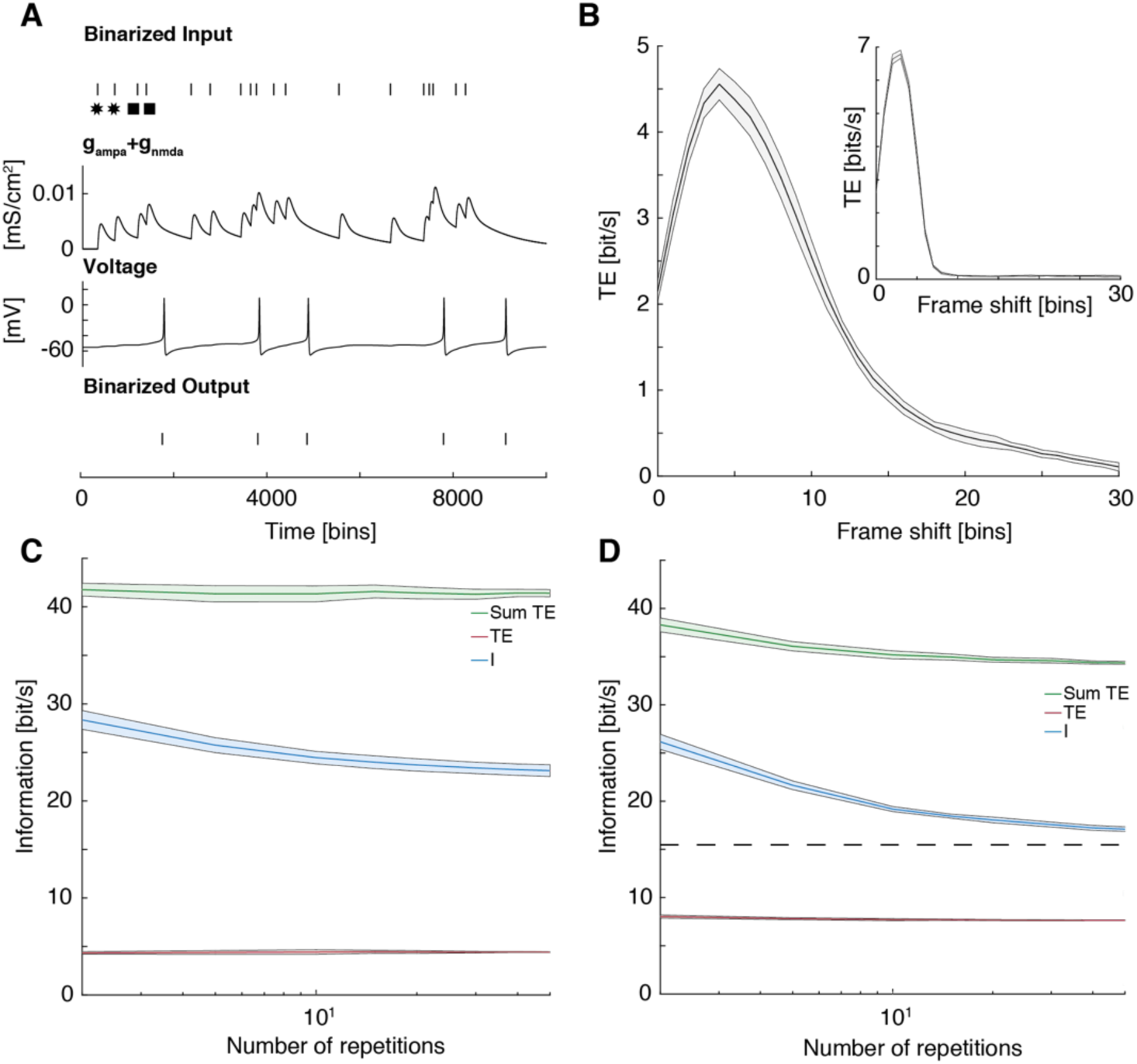
Application to a biophysical Hodgkin-Huxley-type model. (**A**) Poisson spike trains and transmission probabilities are replaced by an experimentally-calibrated Hodgkin-Huxley-type model for thalamic relay cells that recapitulates transmission properties at thalamic relay synapses. 1 time bin = 3 ms. (**B**) Synaptic and neuronal dynamics can lead to output action potentials being generated in different time bins than the incoming input (‘frame shift’). This leads to transfer entropy being positive over multiple temporal frame shifts between the binarized input and output sequences. Mutual information calculated following [11] is immune to that issue. (**C**) Because of this, direct comparison between mutual information (*I*; blue) and transfer entropy (*TE*; calculated using [17]; red: peak value from B; green: integral over *TE* in B) can be difficult. (**D**) This effect can be recapitulated in simple Poisson spike trains by adding gaussian temporal jitter to the timing at which an output spike is generated (see inset in B). *I* is unaffected when the temporal jitter is preserved over repetitions (compare with Figure 2A), while the results obtained for *TE* recapitulate what is observed with the Hodgkin-Huxley-type model (compare with C). In each case, shaded areas indicate the standard error of the mean.

The MATLAB package we used (see Materials & Methods) offers the possibility to compute *TE* with various time frame shifts between the input and output sequences [17]. This is equivalent to simply shifting the frame of reference of the output binary sequence by a set number of bins. Output spikes could be significantly delayed with respect to input spikes for instance in the case of long conduction delays. Here, however, delays are more likely to be due to the specific integration properties of the postsynaptic neuron. Figure 5B shows the evolution of *TE* when plotted versus this shift. It shows that *TE* does not peak at shift 0, but rather raises from about 2bits/s for no frame shift, to about 4.5bits/s at a frame shift of about 15 ms (5 time bins), before decaying again. Note that this is different from what we have observed in a previous study at the cortical synapse between thalamic relay cells and layer 4 spiny stellate cells, where we observed instead that *TE* between the input and output sequences was maximal for no frame shift, and simply decaying for positive shifts [10] (Figure S2 therein). While we have no *a priori* explanation for this finding, this is obviously due to a relatively systematic frame shift between the timing of incoming action potentials and the timing of outgoing action potentials. The rise times of the synaptic conductances and of the membrane potential, i.e. the membrane time constant, might play a role in this observation. By comparison, in the above-mentioned study ([10]), neurons were simulated, or experimentally-recorded, in high-conductance states, which would have shortened their membrane time constant [18]. Here, conductances are only briefly opened after an incoming input action potential. However, this is unlikely to be the full explanation and we hypothesize that this systematic frame shift might also be partially related to the fact that a single action potential will often fail to elicit an output action potential (failed transmission). Instead, it has been known for a while that two incoming action potentials in short succession are usually necessary to trigger an output action potential at those synapses [19], and the input binary sequence we generated based on *in vivo* recordings have non-Poissonian statistics with a preferred interspike interval of ∼10 ms [8] (Figure 4A inset). This effect is apparent in Figure 5A, when two consecutive incoming action potentials separated by 18 ms fail to elicit an output action potential (marked by stars). Immediately after that, two consecutive incoming action potentials separated by 9 ms trigger an output action potential after the second of those has reached the synapse (marked by squares).

In Figure 5C, we plotted the two most obvious measures that can be derived from the curve in Figure 5B, the peak value of *TE* and the integral value of *TE* (calculated over frame shifts from 0 ms to 90 ms) [17]. Both display the same rapid convergence to a stable value, with respect to the size of the dataset, than *TE* in Figures 3 and 4, but they both converge to quite different values (4.4 bits/s and 41.4 bits/s respectively), both different than the value *I* converges to (23.1 bits/s).

In order to test the hypothesis that the strong discrepancy we observed here between matching *I* and *TE* predictions in the simple scenarios graphed in Figures 3 and 4, where action potentials are transmitted from the input to the output sequence in the same corresponding time bin with no temporal frame shift (unless transmission fails), and mismatching *I* and *TE* predictions when using a Hodgkin-Huxley-type model with non-trivial transmission properties (Figure 5A-C), we reproduced the simulations of Figure 3A using simple transmission probabilities and Poissonian inputs. However, this time, we implemented a random shift between every input and output action potentials. This random (positive only) shift followed a Gaussian distribution with mean = 16 ms and standard deviation = 3.7 ms (see Materials & Methods). In the case of *I*, that random shift was systematically reproduced for each pair of inputs and outputs action potentials in each repetition. The inset in Figure 5B shows how the random shift broadens the distribution of TE versus the frame shift. Figure 5D shows that this was sufficient to reproduce the discrepancies described above (compare Figures 3A, 5C and 5D), suggesting that this is indeed the reason behind this observation. Interestingly, by construction, when using the direct method, *I* is immune to that issue (again, compare Figures 3A, 5C and 5D).

Thus, while *TE* offers great stability in its predictions even for relatively small datasets, which will be of interest to experimentalists who might not be able to collect very large datasets due to various technical constraints, the values obtained through that method might not be directly comparable to the mutual information in non-trivial scenarios, in which any kind of systematic frame shift can be expected.

Finally, we wanted to compare the application of these two information theoretic measures to a biophysically relevant scenario. We have recently demonstrated in computational models and experiments that a number of synaptic features can be explained as a trade-off between information flow and energy consumption: the low release probability of weak central synapses [7, 9], the postsynaptic conductance at strong thalamic relay synapses [8], and the postsynaptic conductance at weak cortical synapses [10]. All these studies demonstrate that synapses and neurons, appear to be designed to maximize information flow per energy units (bits/ATP) rather than per time units (bits/sec). More recently, similar results have also been obtained at hippocampal synapses [13]. These results suggest this as a widespread principle in the brain. The existence of an information flow over energy optimum is not surprising. It stems from the basic principle that energy consumption scales roughly linearly with the biophysical parameters we studied (release probability or postsynaptic conductance), while information flow scales sigmoidally with those parameters, because of the necessity to overcome noise [7, 9], or energetic barriers (the threshold for action potential generation) [8, 10]. Thus, information flow over energy, dividing a sigmoid by a linear function, leads to a single well-defined optimum. In particular, we have demonstrated that very mechanism for the postsynaptic conductance at strong thalamic relay synapses, using the same experimentally-calibrated biophysical Hodgkin-Huxley-type model we have used here [8]. Note however that in that original study, we injected in the Hodgkin-Huxley-type model experimentally recorded conductances with a rather limited number of repetitions (N = 5), to estimate *I* with the direct method. Here, using the approach highlighted above to generate *in vivo*-like input spike trains and synaptic conductances (see Figure 5A), we can stimulate that model with any number of repetitions of any length.

We thus proceeded to reproduce here the finding that thalamic relay synapses maximize information flow per energy units (bits/ATP) at experimentally-observed physiological conductances, quantifying information flow using both the mutual information and transfer entropy as above. Specifically, we injected *in vivo*-like conductances generated as in Figure 5A in our experimentally-calibrated biophysical Hodgkin-Huxley-type model of thalamic relay cells. We then varied the postsynaptic conductance by applying a multiplying factor (gain) between 0 and 10 to the injected conductance, like we have done in previous studies [8, 10], the experimentally-observed physiological conductance corresponding to gain = 1. Figure 6A shows information flow across the thalamic relay synapse when modulating the synaptic conductance (gain), quantified as before with *I* (using the direct method) or *TE*. In the latter case, both *TE* measured at its peak frame shift (*TE*_*peak*_; see Figure 5C), or integrated over all frame shifts (*TE*_*sum*_) are reported. As expected from Figure 5C, each measure (*I, TE*_*peak*_ and *TE*_*sum*_) yields different results, but all three grow sigmoidally with the gain. We also additionally quantified the corresponding energy consumption by counting the ATP molecules necessary to fuel the Na,K-ATPase electrogenic pump that restores the ion gradients disturbed by ions flowing through postsynaptic receptors and ion channels (see Methods). Figure 6B shows the energy consumption concomitant to neuronal activity, when accounting only for ionic flows at the modulated postsynaptic conductance, or when accounting for all ionic flows in the postsynaptic neuron (i.e. including also the ions flowing through the voltage-gated channels that underlie action potentials; total energy budget). In both cases, we observed a roughly linear relationship between the modulated postsynaptic conductance and the energy consumption expressed in ATP molecules consumed per second. Note however that in the case of the total energy budget, the relationship is piecewise linear, with a different, smaller slope, for low postsynaptic gains at which no output action potential is generated.

**Figure 6.**
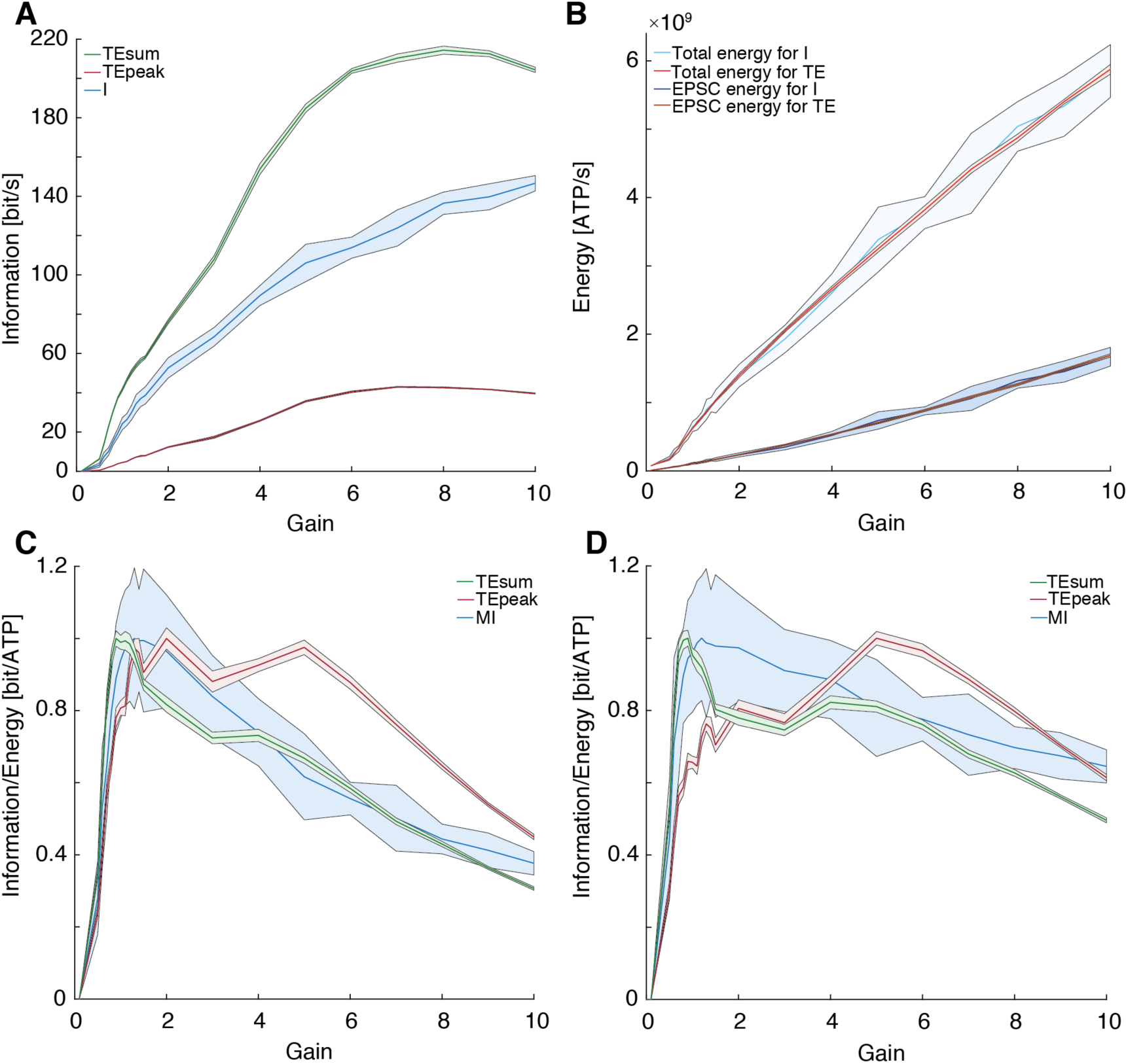
Information flow vs. energy. (**A**) Information transfer at modelled thalamic relay synapses calculated using the Hodgkin-Huxley-type model as a function of the gain of the synapse (normalized synaptic conductance). (**B**) Total energy consumption and EPSC energy consumption used to convey the information calculated in A. (**C**) Normalized ratio between information (A) and EPSC energy (B) as a function of the normalized synaptic conductance. (**D**) Normalized ratio between information (A) and total energy consumption (B) as a function of the normalized synaptic conductance.

We can then evaluate how information flow, quantified using either *I* or *TE*, relates to the concomitant energy consumption at different postsynaptic gains. Figure 6C shows that this relationship (normalized to its peak) has a single well-defined peak close to the physiological gain of the synapse (gain = 1), when quantifying information flow using *I* or *TE*_*sum*_. In both those cases, this ‘energetic efficiency of information flow’ curve peaks close to gain = 1 and closely resembles what has been reported earlier, either for experimental data or in computational models [7-10]. When quantifying the value of information flow using *TE*_*peak*_ however, this relationship appears much broader with no clear discernable peak between gains ∼1 and ∼5. Finally, Figure 6D shows the same results, but using the total energy budget to quantify the energy consumption of the neuron (higher traces in Figure 6B), rather than only the energy consumption imparted by the modulated postsynaptic conductance. The results are however broadly similar to those displayed in Figure 6C. Again, the curves corresponding to *I* and *TE*_*sum*_ match each other and match what has been reported earlier, while the curve corresponding to *TE*_*peak*_ has a broad profile with no clear peak. Therefore, it appears that even though *I* and *TE*_*sum*_ yield different raw values (see Figures 5C and 6A), their predictions can be compared in the current scenario when normalized, unlike *TE*_*peak*_. *TE*_*sum*_ might therefore be a better alternative than *TE*_*peak*_ for comparing to mutual information across studies, although this will have to be systematically verified on a case by case basis.

## Discussion

Here, we set out to evaluate systematically the performance and trade-offs of mutual information and transfer entropy, when applied to binary spike trains in the context of information flowing between individual neurons and across synapses. While these information theoretic measures are popular among theoretical and computational neuroscientists, it can be argued that they have found only limited usage among experimentalists due to the relative complexity in applying them. This is especially true in experimental contexts where acquisition of large datasets is prohibitive. The systematic biases these measures suffer from when applied to limited datasets has led to the development of a number of corrective techniques [6], which, while they improve their performances, do not necessary help in making these techniques more widely accessible. However, it can be argued that more widespread use of information theory will be essential going forward in neuroscience, especially when considering the development of *normative* theories [20] linking information processing to the energetic capacity of the brain [7-10, 21].

Here, we have used a MATLAB package readily available from online repositories [17] to calculate transfer entropy (*TE*) in simple scenarios. We show in these scenarios that *TE* outperforms mutual information when little data is available (Figures 3 and 4) with little need to apply corrective measures. Note that we did alternatively perform the simulations displayed in Figures 3 and 4, but without applying the correction to infinite word lengths (see Figure 2), and that did not change qualitatively the results (not shown). This is however to be expected when using Poisson spike trains (Figure 3), where each input action potential is generated independently from preceding action potentials and no long-range correlations are present in the input sequence [3, 11].

The mutual information is defined as the difference between the two entropies *H*_*total*_ and *H*_*noise*_. For words of length *N, H*_*total*_ calls for the evaluation of *2*^*N*^ independent probabilities from the dataset. *H*_*noise*_ on the other hand calls for the evaluation of *2*^*2N*^ independent probabilities from the same dataset. It is thus common that many of these independent probabilities appearing in the calculation of *H*_*noise*_ will be evaluated to be null because no corresponding event will be observed in a limited dataset. As a consequence, it is common to underevaluate *H*_*noise*_. The same issue is also true for *H*_*total*_, but since far fewer probabilities need to be estimated to calculate *H*_*total*_, it is commonly less underestimated than *H*_*noise*_. The common outcome is then that the mutual information is grossly overestimated for limited datasets. This overestimate decreases in amplitude as the size of the dataset increases (see Figures 3 and 4, and ref. [6] for an excellent discussion of these issues).

Transfer entropy, on the other hand, does not seem to suffer from that problem, at least not in a similar amplitude (Figures 3 and 4). We do not have at this time a definitive explanation as to why that is. Transfer entropy can also be written as the difference between the two entropies H(**X**^-^|**Y**^-^) and H(**X**^-^|Y^+^,**Y**^-^) (see Materials and Methods), but these are two conditional entropies, i.e. they both call for the evaluation of *2*^*2N*^ independent probabilities for words of length *N*. We prudently speculate that this contributes to balancing the systematic errors in evaluating each entropy, and that this leads to a better overall estimation of *TE*, even with limited data (see Figures 3 and 4).

These results suggest that *TE* might be a better choice for experimentalists over *I* due to its lower sensitivity to the size of the dataset, and due to its relative simplicity of use. However, the situation gets more complex when considering a more realistic biophysical scenario. Figure 5 shows that when systematic frame shifts occur between input and output sequences, i.e. when the input and output action potentials do not happen in matching time bins due to transmission delays or due to the integrative properties of the neuron under consideration, the use of *TE* requires a more careful examination of the detailed biophysics at play. It also becomes difficult to directly compare results obtained with *I* and *TE*. In Figure 5, using a Hodgkin-Huxley-type model for thalamic relay cells, we show that the specific integration properties of that modelled neuron lead to transfer entropy being ‘distributed’ over time shifts between the input and output sequences (see Figure 5B). As noted above however, this is not always the case, as we have observed in a previous study (in a different setting) that *TE* between the input and output sequences was maximal for no frame shift, and simply decaying for positive shifts [10]. We additionally provide convincing evidence that this observation is due to a systematic shift by recapitulating the results of Figure 5C using systematically shifted Poisson spike trains. By construction, the mutual information estimated using the ‘direct’ method is immune to that issue.

The fact that *TE* appears to be distributed over time thus poses the question of what feature of *TE* to actually use. The two most obvious features of the curve in Figure 5B are its peak and its integral. Both appear very stable with respect to the size of the dataset but none of them matches the value predicted by mutual information (Figure 5C), making systematic comparisons between these measures difficult. In Figure 6 we apply both features of *TE* (peak and integral) and *I* to the evaluation of the energetic optimality of information transfer at thalamic relay synapses [8]. Our results recapitulate the original experimental and computational finding that those synapses appear to maximize not the information flow (bits/sec), but the ratio of information per concomitant energy use (bits/ATP; see Figure 6). In that context, the integral of *TE* is the feature that matches best the curve obtained using *I*. Both predict the energetically optimal gain of the synapse to be close to 1, the physiological gain of the synapse. The peak of *TE* on the contrary yields markedly different predictions.

These results suggest that a detailed computational study of the system under investigation should be systematically performed prior to applying transfer entropy to experimental data, and that when experimental conditions permit, mutual information estimated using the ‘direct’ method might provide a more straightforward applied measure. In particular, it is important to test for the presence of transmission delays or of integrative properties in the system under investigation that might lead to the kind of effects described here. When it cannot be demonstrated, as we have done in [10], that output action potentials occur mostly in the same time bin than impinging action potentials, it might be best to use mutual information instead.

## Materials & Methods

We are interested in characterizing the properties and performance of two information theoretic measures, mutual information (*I*) and transfer entropy (*TE*), in the context of energy consumption and efficacy of information transfer at individual synapses. In the following, we will apply both these measures to assess information flow from the binary input spike train of a synapse to the binary output spike train generated by the postsynaptic neuron. Below, we start by describing how the test data for this characterization were generated.

### Synthetic spike trains

The first dataset we tested transfer entropy and mutual information on are synthetic Poisson spike trains. In this scenario, the input was a randomly generated Poisson spike train, and the output was created by copying that input spike train, applying for each transmitted spike a certain probability of transmission failure, and for each time bin a certain probability of spontaneous firing, even in the absence of an input spike in the matching input time bin. Unless specified otherwise, all time bins in this manuscript are 3 ms. This simple scenario matches the propagation of individual spikes in the visual pathway, and numerical values for the probabilities were derived from experiments measuring the propagation of spikes at thalamic relay cells in the lateral geniculate nucleus and between thalamic relay cells and layer 4 spiny stellate cells in the primary visual cortex (see numerical values in Table 1 in the Results section above). Unless stated otherwise, each result is the average of 10 independent simulations.

Additionally, in Figure 5, we tested adding a random frame shift between input and output action potentials. We added to the timing of output action potentials a random gaussian delay centered at +16 ms with a standard deviation of ±3.7 ms.

### Input based on biological recordings

The first step towards a more realistic model was to generate inputs similar to those that can be observed impinging onto the cells of interest. To do that, input spike trains were generated according to the distribution of input interspike intervals recorded *in vivo*. The *in vivo* distribution of interspike intervals was used to generate the cumulative distribution function of the intervals, and this function was in turn used as the generative function for the input (see insets in Figure 3). The output was generated as before, with probabilities of failure of transmission and spontaneous firing. 10 simulations were performed, and the mean was taken. Numerical values follow the values given in Table 1.

### Hodgkin-Huxley type model

While using transmission and spontaneous firing probabilities as above is expedient, this can never fully capture the complexity of the biophysical processes that lead to spiking. In order to remediate to that issue, we adapted the single-compartment Hodgkin-Huxley-type model of thalamic relay cells by Harris *et al*. [8]. Details about how that model was carefully calibrated onto experimental data can be found in [8]. Our adjustments are detailed below.

Briefly, the Hodgkin-Huxley-type model of thalamic relay cells in the lateral geniculate nucleus was adapted from earlier models [22-24], and follows the formalism devised by Hodgkin and Huxley with:

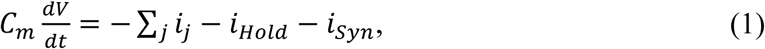

where *C*_*m*_ = 1 µF/cm^2^ is the membrane capacitance, *V* is the membrane voltage (in mV), *i*_*Hold*_ is the injected current, *i*_*Syn*_ is the synaptic current and *i*_*j*_ are the intrinsic currents. The cell surface area was 1.52·10^−4^ cm^2^. All currents and conductances are subsequently reported per unit surface area (cm^2^). Following Bazhenov and colleagues [22], the intrinsic currents included a leak current *i*_*L*_, a potassium leak current *i*_*KL*_, an A-type potassium current *i*_*A*_, a T-type low threshold calcium current *i*_*T*_, an h-current *i*_*h*_, a fast sodium current *i*_*Na*_ and a fast potassium current *i*_*K*_. All the intrinsic currents had the same general form:

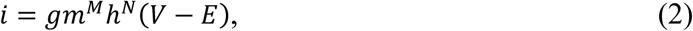

where for each current *i, g* is the maximal conductance, *m(t)* is the activation variable, *h(t)* is the inactivation variable, *E* is the reversal potential and *M* and *N* are the number of independent activation and inactivation gates.

The *i*_*h*_ current was given by:

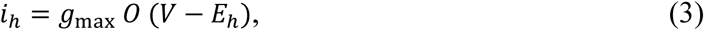

with *E*_*h*_ = –43 mV [25]. *g*_max_ = 0.0254 mS/cm^2^ was set to match experimental data (see [8] for further details). The time dependence of the gating variable *O* was defined by:

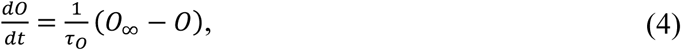

with time constant *τ* _*O*_ = 1/[*e*^(− 14.59−0.086 · *V*)^ + *e*^(− 1.87 + 0.0701 · *V*)^](in ms) and steady-state variable *O*_∞_ = 1/[1 + *e*^((*V* +75)/5.5)^] [25].

The leak currents were given by:

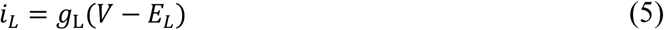

and:

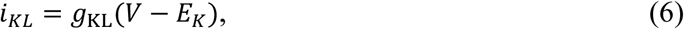

with *E*_*L*_ = –70 mV [24]. *E*_*K*_ was set to match the effective potassium reversal potential used in the experiments in [8]: *E*_*K*_ = –105 mV, while *g*_L_ = 0.025 mS/cm^2^ and g_KL_ = 0.025 mS/cm^2^ were manually adjusted to match both the average input resistance at the resting membrane potential and the resting membrane potential as recorded in experiments.

The A-type potassium current was given by:

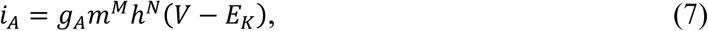

with *M* = 4 and *N* = 1. The time dependence for *m* and *h* was defined as for *O*, with:

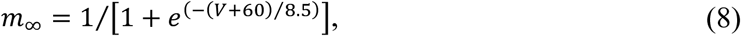

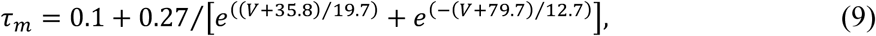

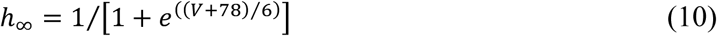

and:

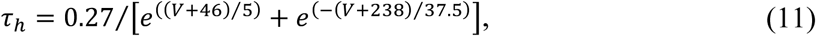

if *V* < –63 mV, and *τ* _*h*_ = 5.1 ms otherwise [22, 23].

The T-type calcium current was given by:

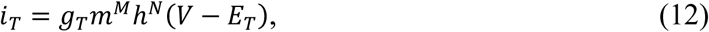

with *M* = 2 and *N* = 1. The time dependence for *m* and *h* was defined as for *O*, with:

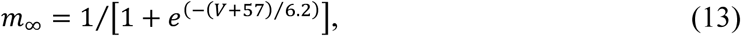

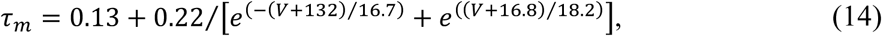

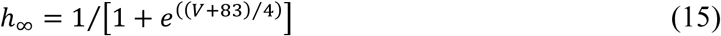

and:

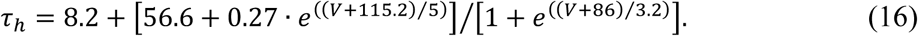

*E*_*T*_ is given by 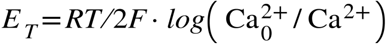 with *F* = 96489 C/mol the Faraday constant, *R* = 8.314 J mol^-1^ K^-1^ the gas constant, T = 309°K the temperature and 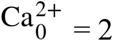 mM the extracellular calcium concentration. The intracellular calcium dynamics were defined by:

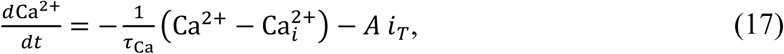

with 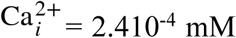 mM, the baseline intracellular calcium concentration, and *A* = 5.1810^−5^ mM cm^2^ ms^-1^ µA^-1^, a constant.

The fast sodium current was defined by:

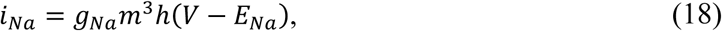

with *E*_*Na*_ = +90 mV. The maximal conductance *g*_*Na*_ = 4.4 mS/cm^2^ was also set to match experimental data [8]. The time dependence for *m* and *h* was defined by:

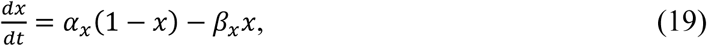

where x stands for either *h* or *m* and with [26]:

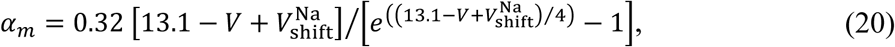

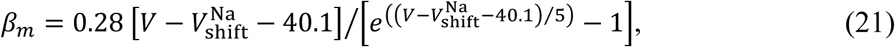

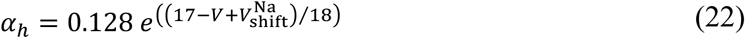

and:

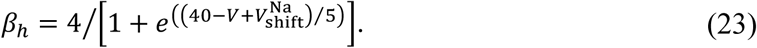

The fast potassium current was given by:

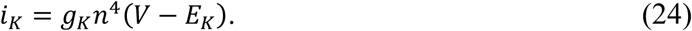

The maximal conductance *g*_*K*_ = 3.3 mS/cm^2^ was set to match experiments (again see [8] for further details). The time dependence for *n* was defined as for the sodium gating variables *m* and *h* with:

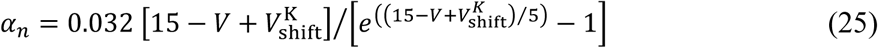

and:

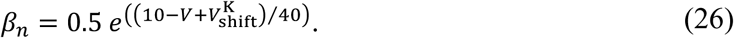

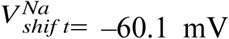 and 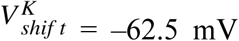 were manually adjusted to allow the model to be depolarized to –55 mV without spontaneously spiking. *i*_*Hold*_ = -2.05 µA/cm^2^ was set in subsequent simulations so as to hold the model at –55 mV. For a cell surface area of 1.5210^−4^ cm^2^, this corresponds to an injected current of ∼310 pA, similar to experimentally measured values of 30-550 pA [8]. *g*_*A*_ = 3 mS/cm^2^ and *g*_*T*_ = 1.8 mS/cm^2^ were set so that the model achieved an output frequency, when stimulated with the synaptic conductance recorded in [5], similar to the average frequency observed in experiments.

### Synaptic currents

Individual synaptic currents experimentally recorded in ref. [8] were fitted by sums of exponentials:

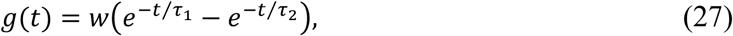

yielding the parameters given in Table 2. The synaptic current is then given by:

**Table 2.**
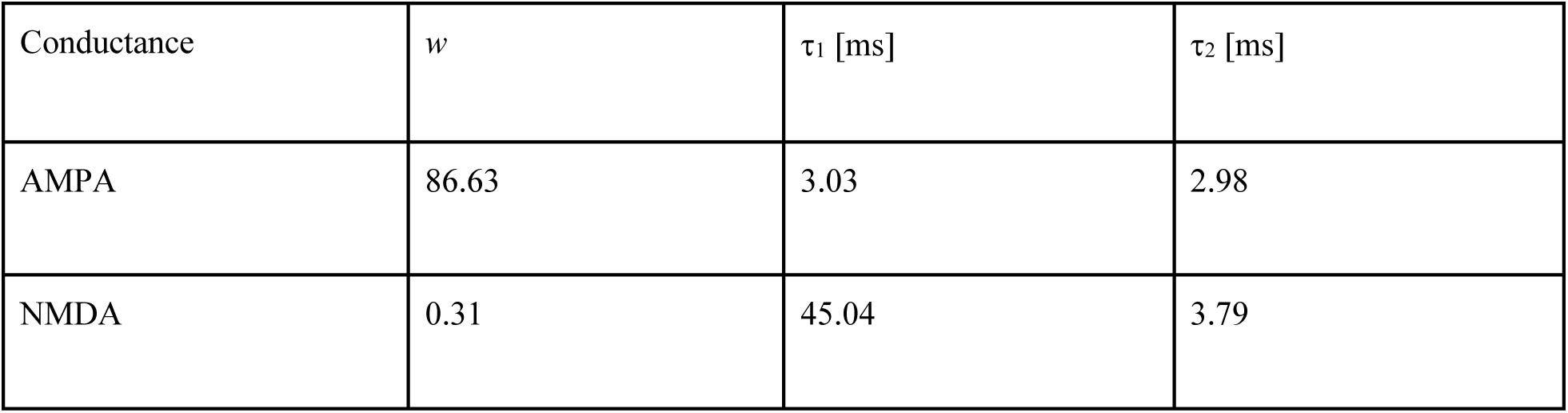
Parameters of individual AMPA and NMDA synaptic conductances.

| Conductance | $w$ | $\tau_1$ [ms] | $\tau_2$ [ms] |
| --- | --- | --- | --- |
| AMPA | 86.63 | 3.03 | 2.98 |
| NMDA | 0.31 | 45.04 | 3.79 |

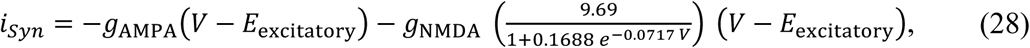

with *E*_excitatory_ = 0 mV, and where *g*_AMPA_ and *g*_NMDA_ describe the time courses of individual AMPA and NMDA synaptic currents.

### Information theory

For two coupled physical systems that produce realizations *x* and *y* of random variables *X* and *Y*, the mutual information *I* is defined by [27]:

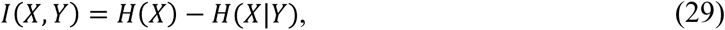

where *H*(*X*) = ∑_*x*_ *p*(*x*) is the Shannon entropy for the probability distribution *p(x) = p(X=x)* of the outcome *x* of the random variable *X* and *H*(*X*|*Y*) = ∑_*y*_ *p*(*y*) ∑_*x*_ *p*(*x*|*y*) log_2_-(1/*p*(*x*|*y*)) is the conditional entropy. The mutual information can then be written as:

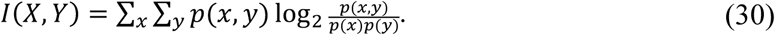

Because *H(X)* is the total average information in the variable *X* and *H(X*|*Y)* is the average information that is unique to *X*, the mutual information represents the shared information between the two processes or, in other words, the deviation from independence of the two processes. Equations (29) and (30) are symmetric under the exchange of the two variables and thus do not contain any directional indication.

The transfer entropy TE of the same random variables is defined as [27]:

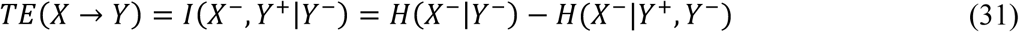

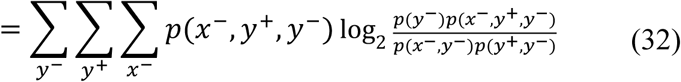

where ***X***^*-*^ and ***Y***^*-*^ denote the past state of the two processes with outcomes *x*^*-*^ and *y*^*-*^, and *Y*^*+*^ is the future random variable of *Y* with outcome *y*^*-*^.

The transfer entropy represents the amount of predictive information actually transferred from process *X* to process *Y*. As can be seen in Equation (31), the transfer entropy is not usually symmetric under the exchange of *X* and *Y*.

### Equivalence between I and TE

In a scenario where ***X***^*-*^ and *Y*^*+*^ are both independent of ***Y***^*-*^ (see Equation (31) above), which is for instance realized in the scenarios displayed in Figure 3, the transfer entropy *TE* can be written as:

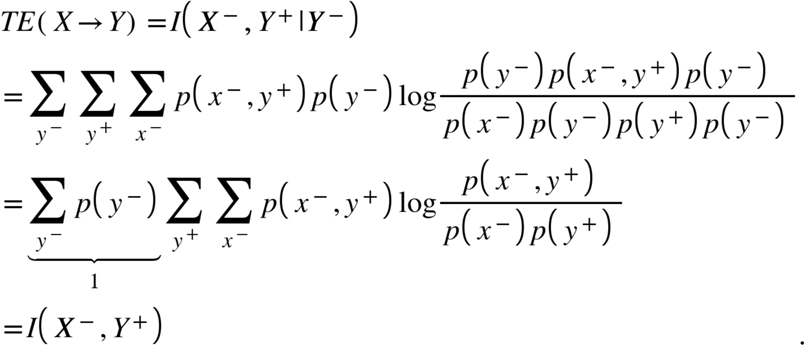

In this simplified case, *TE* is equivalent to *I*. In Figure 3, we have used this fact, together with the fact that with knowledge of all the probabilities entering in Equations (29)-(30), we can directly calculate – rather than estimate from the data – the theoretical value of *TE = I*.

### Estimation methods

Poisson spike trains are already binary. The output spike trains generated with the Hodgkin-Huxley-type model were binarized using a time bin of 3 ms using the function *findpeaks* in MATLAB (The Mathworks, Natick MA).

Mutual information was evaluated using the so-called ‘direct’ method devised by Strong and colleagues [11]. As noted in the Results section, we only applied the second of the two corrections from the original method, i.e. the extrapolation to infinite word lengths (see Figure 2, Results and ref. [8] for further details).

Transfer entropy was calculated using the MATLAB package by Ito and colleagues [17], following similar procedures as in [10]. In particular, a baseline value *TE*_*noise*_ was calculated by randomly shuffling words in the output and calculating the transfer entropy between the input and the shuffled output. The transfer entropy values reported in the manuscript are *TE = TE*_*raw*_ *-TE*_*noise*_ after extrapolation to words of infinite lengths similar to the correction applied for calculations of the mutual information.

Repetitions of 128 s were used for calculations of the mutual information using the ‘direct’ method. Each data point is the average of N = 10 independent simulations. For direct comparison, we used datasets of the same length for calculations of the transfer entropy. For instance, if we had 50 repetitions of 128 s for the ‘direct’ method (6400 s total), we used a single input spike train of 6400 s for calculation of the transfer entropy.

### Information and energy

To calculate information transfer at the simulated synapse, output spike trains were processed as detailed above. To calculate the metabolic cost incurred by the modelled cell, the Na^+^ component of *i*_*Syn*_ was integrated and converted to the corresponding ATP consumption per unit time, while the same procedure was followed for *i*_*Na*_, the Na^+^ component of *i*_*h*_ and Ca^2+^ entry via *i*_*T*_, and this was added to the ATP used on *i*_*Syn*_. For *i*_*Syn*_, the conductance was scaled by 7/13 (derived from the reversal potentials *E*_excitatory_ = 0 mV, *E*_*Na*_ = +90 mV and *E*_*K*_ = -105 mV) and multiplied by *V -E*_*Na*_ to calculate the contribution of sodium ions. For *i*_*h*_, the conductance was scaled by (*E*_*k*_ − *E*_*h*_)/(*E*_*k*_ − *E*_*Na*_) and multiplied by *V - E*_*Na*_ to isolate the contribution of sodium ions. For *i*_*T*_, we assumed that each calcium ion is exchanged for 3 sodium ions [28]. For each gain in Figure 6, N = 10 simulations were performed, and the mean was taken.

### Simulations

Simulations were run using custom-written MATLAB scripts (The Mathworks, Natick MA). Differential equations were integrated using the built-in solver *ode15s* with an integration time step dt = 0.05 ms. All results presented are the mean of N = 10 independent simulations.

## References

1. Cobb M. The Idea of the Brain: A History: Profile Books; 2020. 480 p.

2. Shannon CE. A mathematical theory of communication. Bell Syst Tech J. 1948;27:379–423.

3. Reinagel P, Reid RC. Temporal coding of visual information in the thalamus. J Neurosci. 2000;20(14):5392–400. PubMed PMID: 10884324.

4. Dayan P, Abbott LF. Theoretical Neuroscience: Computational and Mathematical Modeling of Neural Systems. Cambridge MA: MIT Press; 2001. 576 p.

5. London M, Schreibman A, Hausser M, Larkum ME, Segev I. The information efficacy of a synapse. Nat Neurosci. 2002;5(4):332–40. doi: 10.1038/nn826. PubMed PMID: 11896396.

6. Panzeri S, Senatore R, Montemurro MA, Petersen RS. Correcting for the sampling bias problem in spike train information measures. J Neurophysiol. 2007;98(3):1064–72. doi: 10.1152/jn.00559.2007. PubMed PMID: 17615128.

7. Harris JJ, Jolivet R, Attwell D. Synaptic Energy Use and Supply. Neuron. 2012;75(5):762–77. doi: 10.1016/j.neuron.2012.08.019. PubMed PMID: WOS:000308684300005.

8. Harris JJ, Jolivet R, Engl E, Attwell D. Energy-Efficient Information Transfer by Visual Pathway Synapses. Curr Biol. 2015;25(24):3151–60. doi: 10.1016/j.cub.2015.10.063. PubMed PMID: WOS:000367233400014.

9. Conrad M, Engl E, Jolivet R. Energy use constrains brain information processing. Technical Digest - International Electron Devices Meeting. 2018:11.3.1-.3.3.

10. Harris JJ, Engl E, Attwell D, Jolivet RB. Energy-efficient information transfer at thalamocortical synapses. PLoS Comput Biol. 2019;15(8):e1007226. doi: 10.1371/journal.pcbi.1007226.

11. Strong SP, Koberle R, de Ruyter van Stevenick R, Bialek W. Entropy and Information in Neural Spike Trains. Physical review letters. 1998;80(1):197–200.

12. Schreiber T. Measuring information transfer. Physical review letters. 2000;85(2):461-4. Epub 2000/09/16. doi: 10.1103/PhysRevLett.85.461. PubMed PMID: 10991308.

13. Mahajan G, Nadkarni S. Local design principles at hippocampal synapses revealed by an energy-information trade-off. bioRxiv. 2019. doi: 10.1101/748400.

14. Kostal L, Shinomoto S. Efficient information transfer by Poisson neurons. Math Biosci Eng. 2016;13(3):509-20. Epub 2016/04/24. doi: 10.3934/mbe.2016004. PubMed PMID: 27106184.

15. Kostal L, Lansky P, McDonnell MD. Metabolic cost of neuronal information in an empirical stimulus-response model. Biol Cybern. 2013;107(3):355-65. Epub 2013/03/08. doi: 10.1007/s00422-013-0554-6. PubMed PMID: 23467914.

16. Kostal L, Lansky P. Information capacity and its approximations under metabolic cost in a simple homogeneous population of neurons. Biosystems. 2013;112(3):265-75. Epub 2013/04/09. doi: 10.1016/j.biosystems.2013.03.019. PubMed PMID: 23562831.

17. Ito S, Hansen ME, Heiland R, Lumsdaine A, Litke AM, Beggs JM. Extending transfer entropy improves identification of effective connectivity in a spiking cortical network model. PLoS One. 2011;6(11):e27431. Epub 2011/11/22. doi: 10.1371/journal.pone.0027431. PubMed PMID: 22102894; PubMed Central PMCID: PMCPMC3216957.

18. Destexhe A, Rudolph M, Paré D. The high-conductance state of neocortical neurons in vivo. Nature Reviews Neuroscience. 2003;4:739–51.

19. Carandini M, Horton JC, Sincich LC. Thalamic filtering of retinal spike trains by postsynaptic summation. J Vis. 2007;7(14):20 1-11. Epub 2008/01/26. doi: 10.1167/7.14.20. PubMed PMID: 18217815; PubMed Central PMCID: PMCPMC2901808.

20. Levenstein D, Alvarez VA, Amarasingham A, Azab H, Gerkin RC, Hasenstaub A, et al. On the role of theory and modeling in neuroscience. arXiv. 2020. doi: https://arxiv.org/abs/2003.13825.

21. Aiello LC, Wheeler P. The Expensive-Tissue Hypothesis. Current Anthropology. 1995;36(2):199–221.

22. Bazhenov M, Timofeev I, Steriade M, Sejnowski TJ. Cellular and network models for intrathalamic augmenting responses during 10-Hz stimulation. Journal of Neurophysiology. 1998;79(5):2730-48. PubMed PMID: 9582241.

23. Bazhenov M, Timofeev I, Steriade M, Sejnowski TJ. Computational models of thalamocortical augmenting responses. Journal of Neuroscience. 1998;18(16):6444-65. PubMed PMID: 9698334.

24. McCormick DA, Huguenard JR. A Model of the Electrophysiological Properties of Thalamocortical Relay Neurons. Journal of Neurophysiology. 1992;68(4):1384-400. PubMed PMID: ISI:A1992JV73300034.

25. Huguenard JR, McCormick DA. Simulation of the currents involved in rhythmic oscillations in thalamic relay neurons. Journal of Neurophysiology. 1992;68(4):1373-83. PubMed PMID: 1279135.

26. Traub RD, Miles D. Neuronal Networks of the Hippocampus. Cambridge: CUP; 1991.

27. Wibral M, Vicente R, Lindner M. Transfer entropy in neuroscience. In: Wibral M, Vicente R, Lizier JT, editors. Directed Information Measures in Neuroscience: Springer; 2014. p. 3–36.

28. Attwell D, Laughlin SB. An energy budget for signaling in the grey matter of the brain. Journal of Cerebral Blood Flow and Metabolism. 2001;21:1133–45.

